# Endocrine pancreas-specific *Gclc* gene deletion causes a severe diabetes phenotype

**DOI:** 10.1101/2023.06.13.544855

**Authors:** Emily A. Davidson, Ying Chen, Surendra Singh, David J. Orlicky, Brian Thompson, Yewei Wang, Georgia Charkoftaki, Tristan A. Furnary, Rebecca L. Cardone, Richard G. Kibbey, Colin T. Shearn, Daniel W. Nebert, David C. Thompson, Vasilis Vasiliou

## Abstract

Reduced glutathione (GSH) is an abundant antioxidant that regulates intracellular redox homeostasis by scavenging reactive oxygen species (ROS). Glutamate-cysteine ligase catalytic (GCLC) subunit is the rate-limiting step in GSH biosynthesis. Using the *Pax6-Cre* driver mouse line, we deleted expression of the *Gclc* gene in all pancreatic endocrine progenitor cells. Intriguingly, *Gclc* knockout (KO) mice, following weaning, exhibited an age-related, progressive diabetes phenotype, manifested as strikingly increased blood glucose and decreased plasma insulin levels. This severe diabetes trait is preceded by pathologic changes in islet of weanling mice. *Gclc* KO weanlings showed progressive abnormalities in pancreatic morphology including: islet-specific cellular vacuolization, decreased islet-cell mass, and alterations in islet hormone expression. Islets from newly-weaned mice displayed impaired glucose-stimulated insulin secretion, decreased insulin hormone gene expression, oxidative stress, and increased markers of cellular senescence. Our results suggest that GSH biosynthesis is essential for normal development of the mouse pancreatic islet, and that protection from oxidative stress-induced cellular senescence might prevent abnormal islet-cell damage during embryogenesis.

## Introduction

Reduced glutathione (GSH) is the most abundant intracellular non-protein thiol and, arguably, the most important intracellular antioxidant in all vertebrates. GSH is biosynthesized in the cytosol of most mammalian cells in a series of ATP-dependent enzymatic reactions, limited by the activity of glutamate-cysteine ligase (GCL) [1]. GCL is a heterodimeric holoenzyme whose catalytic subunit, GCLC, is required for GSH biosynthesis. GSH is responsible for maintaining the cellular redox environment, largely by directly scavenging reactive oxygen species (ROS), acting as a cofactor for antioxidant enzymes, and/or participating in redox signaling events [2]. Depletion of GSH induces oxidative stress, a pathophysiological state in which ROS production and antioxidant capacity are imbalanced in favor of an oxidative state. Oxidative stress is associated with perturbed cellular function [3–5] and structure [6, 7] which can progress to cell death [8–10].

The importance of GCLC expression (and subsequent GSH synthesis) in physiology has been demonstrated in mouse models. For example, whole body deletion of *Gclc* is embryonic lethal [11], hepatocyte-specific *Gclc* expression is essential for liver function [12], *Gclc* deficiency in the central nervous system promotes neurodegeneration [13], and T cell function is disrupted by *Gclc* deletion [14]. Recently, we reported that deletion of *Gclc* in the developing eye caused microphthalmia (a disease involving abnormally small eyes) [15].

In this model, the Cre/*loxP* recombination approach was used to delete *Gclc* and the expression of Cre was directed to the developing eye using *Le-Cre* transgenic mice. Herein, Cre expression is activated by the P0 promoter of the *Pax6* gene and expressed in surface ectoderm-derived ocular tissues from embryonic day ∼9.5 [16]. In addition to possessing a crucial role in eye development [16], *Pax6* is a transcription factor essential for regulating pancreatic islet development [17]. Therefore, *Le-Cre* transgenic mice express Cre in endocrine progenitor cells from embryonic day ∼9.5 [18].

During characterization of the microphthalmia phenotype, *Gclc* knockout (KO) mice were unable to survive past 60 days of age. Furthermore, the *Gclc* KO mice had progressively poor health. By general observation, *Gclc* KO mice had increased urination (indicated by wet bedding and a pungent smell) at ∼35 days of age and beyond. Additionally, *Gclc* KO mice displayed a hunched posture and visible weight loss after ∼40 days of age. When quantified (**S1a**), *Gclc* KO mice weighed significantly less than *Gclc* CTRL mice at 42 days of age. This weight loss was rapid, between 40 and 42 days of age*, Gclc* KO mice lost ∼30% of bodyweight. This significant weight loss was deemed a humane concern and mice required euthanasia. Clinically, these symptoms mimicked those observed in uncontrolled diabetes. Prior to required euthanasia, we measured fasting blood glucose levels in *Gclc* KO mice and, compared to *Gclc* CTRL mice, *Gclc* KO mice displayed significantly higher fasting blood glucose levels (**S2a**). While our original research goal was to delete *Gclc* in the developing eye, we inadvertently deleted *Gclc* in the developing endocrine pancreas, which is in line with the reported Pax6-drive CRE expression [18], leading to a severe diabetes phenotype.

In this study, we characterized the nature of the diabetic phenotype observed in the *Gclc* knockout (KO) mice, and determined that deletion of *Gclc* in endocrine progenitor cells causes an age-related, progressive diabetes phenotype. Assessment of glucose homeostasis revealed *Gclc* KO mice displayed post-weaning (≥ 25 days of age) increased fasting blood glucose, decreased fasting blood insulin levels, and impaired glucose tolerance. Changes to glucose homeostasis were accompanied by islet-specific progressive pathological changes manifesting as larger islets (at postnatal day (P) 21), cytoplasmic vacuolization and diffuse islet hormone expression (at P30 – P35), and reduced islet size and islet hormone expression (at P40). Prior to the onset of overt diabetes, newly weaned (P21) *Gclc* KO mouse islets (although enlarged) displayed reduced glucose stimulated insulin secretion. Further molecular characterization revealed, *Gclc* KO mouse islets had decreased islet hormone expression, increased markers of oxidative stress, and increased markers of early senescence. Collectively, we demonstrate that GSH biosynthesis is essential for normal development of the mouse pancreatic islet. Our results suggest that, protection against oxidative stress-related cellular senescence may underly the essentiality of GSH to pancreatic islet development.

## Experimental Procedures

### Animals

*Le-Cre (Tg(Pax6-cre,GFP)1Pgr)* transgenic and *Gclc^f/f^* mice were generated as previously described [12, 16]. *Le-Cre* transgenic mice were originally produced by Dr. Ruth Ashery-Padan (Department of Human Molecular Genetics and Biochemistry, Sackler Faculty of Medicine, Tel Aviv University), and were obtained from Dr. David Beebe (Department of Ophthalmology and Visual Sciences, Washington University School of Medicine). *Gclc^wt/f^*/*Le-Cre^Tg/-^* male mice were continuously crossed with *Gclc^f/f^* female mice to generate two experimental mouse genotypes, i.e., *Gclc^f/f^*/*Le-Cre^−/−^* (*Gclc* CTRL) and *Gclc^f/f^/Le-Cre^Tg/-^* (*Gclc* KO). Mice used in the present study were generated from B6/FVB mixed genetic background. All studies (except those involving P14 mice) were conducted in male mice weaned at P21. For studies conducted in P14 mice, both sexes were used. Mice were provided water and food *ad libitum* and maintained under a 12-hour light/dark cycle. All animal studies were performed in strict accordance with the National Institutes of Health guidelines, under protocols approved by the Institutional Animal Care and Use Committee of Yale University.

### Genotyping

Genomic DNA (obtained from a 2mm ear punch) was extracted using DirectPCR Lysis Reagent (Mouse tail) (Viagen Biotech, Los Angeles, CA) and PCR-grade proteinase K (Roche #RPROTK-RO, Basel, Switzerland) according to the manufacturer’s protocol. Briefly, the ear punch sample was placed in 100μL DirectPCR Lysis Reagent containing 1μL PCR-grade proteinase K. The sample was incubated overnight at 55°C, and subsequently incubated at 85°C for 50 min. Separate PCR reactions were required to genotype each mouse: one detected the *Gclc(wt)* and *Cre(Tg)* alleles, and the other detected the *Gclc(f)*. *Gclc* alleles were determined using primers for *5^’^-CTATAATGTCCTGCACTGGG*, *5^’^-TAGTGAACGCTGTTAAAGG,* and *5′-CGGGTGTTGGGTCGTTTGT*. The presence of the *Cre* transgene was identified using primers for *5^’^-GCGGTCTGGCAGTAAAAACTATC* and *5′-GTGAAACAGCATTGCCACTT*, as described previously [12, 15]. Primers were synthesized by the Yale Keck Oligonucleotide Synthesis facility.

### Measurement of Fasting Blood Glucose and Fasting Plasma Insulin

Fasting blood glucose and plasma insulin measurements were conducted in mice at ages P14, P21, P25, P30, P35, and P40. Prior to blood collection, mice were fasted for 6 h (morning fast, start time of ∼8:00am). For fasting blood glucose measurements, a blood sample was collected from the tail vein and quantified using a blood glucose meter (Bayer Contour^®^, Ascensia Diabetes Care, Basel, Switzerland). For fasting plasma insulin measurement, mice were euthanized *via* the open-drop isoflurane method, and blood was collected from the vena cava. In P14 mice, blood was collected from a small incision in the vena cava using a capillary tube (Hemato-Clad™, Drummond Scientific Company, #1-00-7500-HC/5, Broomall, PA). Each blood sample was immediately placed into a heparin/lithium-coated tube (Beckman Coulter #652825, Indianapolis, IN) placed on ice. To obtain plasma, the blood sample was subjected to centrifugation (11,800 x g) for 1 min at room temperature (RT). For mice older than P14 mice, blood was collected using a 25-gauge needle and immediately placed into a K3EDTA tube (MiniCollect^®^, Greiner Bio-One, Monroe, NC, USA). To obtain plasma, the blood sample was subjected to centrifugation (2,000g) for 10 min at 4°C. The resultant supernatant was isolated and stored at −80°C for latter analysis. Insulin was quantified using ELISA (Mouse Ultrasensitive Insulin ELISA, ALPCO Diagnostics, Salem, NH) according to manufacturer’s instructions.

### Islet Isolation

Islets were isolated from pancreata of P21 mice as previously described [19]. Briefly, mice were euthanized *via* the open drop isoflurane method. The pancreas was removed *en bloc via* a midline incision and placed in ice-cold modified Hanks’ Balanced Salt Solution (HBSS) (Gibco™, Thermo Fisher Scientific, Waltham, MA) containing 3.25% HEPES (Gibco™, Thermo Fisher Scientific, Waltham, MA), 1% 10,000 U/mL penicillin and 10,000 U/mL streptomycin (Gibco™, Thermo Fisher Scientific, Waltham, MA). Pancreata (from 3-5 mice per genotype) were pooled, rinsed with cold modified HBSS, and digested in modified HBSS containing 2mg/mL collagenase P (Roche #11215809103, Basel, Switzerland) and 2% bovine serum albumin (BSA) (Sigma Aldrich, St. Louis, MO). The solution was gently agitated for 2-3 min time periods in a 37°C water bath until the tissue was well-digested (∼4-6 min total digestion time). Islets were purified from other pancreatic tissue using density gradient centrifugation. The digested material was centrifuged at RT, 1,200 RPM for 20 min in Histopaque 1100 solution, containing 45% Histopaque® 1077 (Sigma-Aldrich #10771, St. Louis, MO) and 55% Histopaque 1119® (Sigma-Aldrich #11191, St. Louis, MO, USA). The purified islet-containing supernatant was removed, diluted with ice-cold modified HBSS and centrifuged (1,500 RPM) for 5 min at RT. Supernatant was removed from the pelleted islets and islets were wash with ice-cold modified HBSS twice at 1,000 RPM for 2 min. After the final centrifugation, the supernatant was discarded and the pelleted islets were re-suspended in 4mL of complete RPMI-1640 medium containing 10% fetal bovine serum (Gibco™, Thermo Fisher Scientific, Waltham, MA), 1% HEPES, and 1% 10,000 U/mL penicillin and 10,000 U/mL streptomycin. The suspension was deposited into a 60mm petri dish (Falcon #351007, Columbus, Indiana, USA) and islets were hand-picked from any residual non-islet tissue using a 20uL pipette under a light microscope (Leica DMIL LED, Leica, Wetzlar, Germany) and placed in 4mL of the RPMI-1640 medium in a 60mm petri dish (Falcon #351007, Columbus, Indiana, USA). Isolated islets (∼200-400 islets/4 mL medium) were cultured overnight in a humidified (95%) atmosphere of 5% CO_2_ in air maintained at 37°C before initiation of experiments.

### Western Blot

Islets were isolated, as described above. Islets were hand-picked from the petri dish using a 20uL pipette under a light microscope, placed in a 1.5mL microcentrifuge tube (Eppendorf, Enfield, CT, USA) containing 1mL of RMPI-1640 media, and centrifuged (1,500 RPM) for 3 min at 4°C. The resultant pellet was resuspended in 1mL ice-cold 1x Dulbecco’s Phosphate-Buffered Saline (DPBS) (Gibco™, Thermo Fisher Scientific, Waltham, MA) and subjected to centrifugation (1,500 RPM) for 5 min at 4°C. The supernatant was removed, flash frozen in liquid nitrogen and stored at −80°C for latter protein extraction. The pellet (containing islets) was resuspended in 30uL 1x RIPA lysis buffer (Millipore, Burlington, MA) containing protease inhibitor (cOmplete™, Roche #11836170001, Basel, Switzerland) (1 tablet/10mL) and allowed to incubate on ice for 30 min. After incubation, the suspension was vortexed for 30 sec, sonicated (Branson 5800 Cleaner, Emerson, St. Louis, MO, USA) at 40 kHz for 5 min, and subjected to centrifugation (14,000 RPM) for 10 minutes at 4°C. The supernatant (islet sample) was collected and the total protein was quantified using the Pierce BCA Protein Assay Kit (Thermo Fisher Scientific, Waltham, MA) according to the manufacturer’s instructions. Twenty µg (islet sample) or 2 µg (mouse liver sample: positive control) was mixed with 10uL of Laemmli Sample Buffer (Bio-Rad, Hercules, CA) and denatured at 95°C for 10 min. 40uL of protein sample solution, containing 20µg denatured protein was applied to a 4–20% SDS-PAGE gradient gel (Bio-Rad, Hercules, CA) electrophoresis and semi-dry transferred to a 0.2μm nitrocellulose membrane (Bio-Rad, Hercules, CA) using the Trans-Blot^®^ Turbo™ Transfer System (Bio-Rad #1704150, Hercules, CA). The membrane was blocked using 5% non-fat dry milk (AmericanBio #AB10109, Natick, MA) in 1x Tris Buffered Saline containing 0.1% Tween 20 (TBST) (AmericanBio #AB14330, Canton, MA) for 1 h at RT on an orbital shaker (95 RPM). After blocking, the membrane was incubated overnight in primary antibody (see below) at 4°C on an orbital shaker (100 RPM). Membranes were rinsed three times at RT in TBST for 10 min, and subsequently incubated with HRO-conjugated secondary antibody at RT for 1 h on an orbital shaker (95 RPM). The membrane was then washed three times in TBST for 10 min at RT. Immunolabeled proteins were detected by application of enhanced chemiluminescence Western blotting reagents (Clarity Western ECL Substrate, Bio-Rad, Hercules, CA) according to the manufacturer’s instructions. Immunoblots were visualized using the ChemiDoc Touch Gel Imaging System (Bio-Rad #1708370, Hercules, CA). Primary antibody for rabbit monoclonal anti-GCLC (Abcam #ab207777, Cambridge UK) was used at a dilution of 1:500. The secondary antibody (HRP-conjugated goat anti-rabbit) (Cell Signaling Technology #7074, Danvers, MA) was used at a dilution of 1:1000. Primary and secondary antibodies were diluted in 3% BSA in TBST. Between the probing of each protein target, the membrane was stripped using Restore™ PLUS Western Blot Stripping Buffer (Thermo Fisher Scientific #46430, Waltham, MA) per manufacturer’s instructions. After stripping, the membrane was rinsed three times at RT in TBST for 10 min, blocked, and probed as described above. Total protein, per lane, was determined by Ponceaus S stain (Sigma-Aldrich #P7170, St. Louis, MO), according to manufacturer’s instructions. Target protein band density was normalized to total protein. Analysis was performed in triplicate (n = 3 bands/genotype). Each band represents a pool of ∼400 islets from three independent islet isolation experiments. Band densities were quantitated using NIH Image J software [20, 21]. Data are presented as the mean density (and associated standard deviation) of the normalized total protein in arbitrary units (a.u.).

### Histological and Immunohistochemical Analysis

Histological and immunohistochemical analysis was conducted in mice at postnatal days (P) 21, 25, 30, 35, and 40. For P14 measurements, male and female mice were assessed. For ≥P21 measurements, male-only mice were assessed. Mice were weaned at P21. Mice were euthanized via the open-drop isoflurane method The abdominal cavity was accessed through a midline incision, whole-pancreas removed, and fixed in 10% Neutral buffered formalin at 4°C for 24 hours. After fixation, the tissue was transferred to a 70% ethanol solution and delivered to Yale Pathology Tissue Services for paraffin-embedding, sectioning, and Hematoxylin & Eosin (H&E) staining.

Immunohistochemical staining for insulin and glucagon was conducted using a Tyramide Signal Amplification (TSA) Plus Biotin kit according to the manufacturer’s protocol (NEL700A001KT, Akoya Biosciences, Marlborough, MA, USA), as previously described [15]. Briefly, deparaffinized (immersion in xylene and ethanol) and rehydrated (immersion in water) pancreas sections were heated in boiling 0.01 M citrate buffer (pH 6.6) for 20 minutes for antigen retrieval. After cooling down to RT, tissue sections were immersed in 3% hydrogen peroxide in 30% methanol for 10 minutes at RT to deplete endogenous peroxidase activity. After 4 washes with distilled water, the sections were incubated in Tris-NaCl (TNB) blocking buffer (Akoya Biosciences #FP1020, Marlborough, MA, USA) in a humidified chamber at RT for 1 hour to block nonspecific staining. The sections were then incubated with primary antibodies (see below) at RT for 2 hours. The sections were washed 3 times with PBST (PBS + 0.5% Tween-20) and incubated with HRP-conjugated secondary antibody at RT for 1 hour. The sections were then incubated for 30 minutes in streptavidin HRP (Akoya Biosciences #TS-000300, Marlborough, MA, USA), washed in PBST, incubated for 10 minutes with biotinyl tyramide (Akoya Biosciences #TS-000109, Marlborough, MA, USA), and washed in PBST. The sections were visualized by incubating with 3,3’-diaminobenzidine (DAB) substrate (Vector Laboratories #SK-4100, Newark, CA, USA) solution for 10 minutes and counterstained with hematoxylin (1:10 dilution in distilled water) for 2 minutes. Lastly, the sections were dehydrated in ascending concentrations of ethanol and cover slipped with Permount (#SP15-500, Fischer Chemical, Hampton, NH, USA). Primary antibodies were used at dilutions of 1:200 for Insulin (Cell Signaling Technology #4590S, Danvers, MA, USA) and Glucagon (Cell Signaling Technology #2760S, Danvers, MA, USA), and 1:500 for F4/80 (Cell Signaling Technology #70776 Danvers, MA, USA). The secondary antibody was HRP-conjugated goat anti-rabbit (Cell Signaling Technology #7074, Danvers, MA, USA) at 1:500. Primary and secondary antibodies were diluted in TNB blocking buffer. Staining for neutrophils, T cells, and B cells was conducted by Yale Pathology Tissue Services. Images (40-200x) were collected using an Olympus BX51 brightfield microscope.

### Measurement of Relative Islet Number, Mass, and Size

To quantitate islet number, mass and size, one pancreas section per two H&E-stained slides, separated by at least 100uM, were imaged per mouse pancreata. The section was scanned (at 10x magnification) and digitized using the All-in-One Fluorescence Microscope (Keyence #BZ-X800, Keyence, Osaka, Japan). Individual images were uploaded to ImageJ [22] and manually annotated for total pancreas area and total individual islet area in pixels per pancreas section. Total islet number per mouse ranged from ∼65-124, with variation due to section size. Total pancreas area, defined as the total pixels (the sum of islet and exocrine pancreas pixels), was used to normalize for differences in section size. Calculations were conducted from each pancreas section as follows: (i) relative islet number = (sum of the number of islets/sum of total pixels), (ii) relative islet mass = (sum of total islet pixels/sum of total pixels), and (iii) relative islet size = (average of total islet pixels/sum of total pixels). Measurements were conducted in triplicate, i.e., pancreata from three mice were analyzed per genotype.

### Static Glucose Stimulated Insulin Secretion

Static Glucose Stimulated Insulin Secretion assays (GSIS) were conducted on islets, isolated from mice at P21, using standardized protocols from the Yale University Islet Isolation and Function Core. In brief, islets were isolated, as described above. Isolated islets were seeded at a density of 10-15 islets/well in a HTS Transwell® 96-well permeable supports with receiver plate (Corning #3382 and #3387, Corning, NY, USA) containing islet culture media. To remove the culture media, the plate was centrifuged at RT, 1,000 RPM for 1 minute. The culture media was replaced with 2.5 mM glucose-containing buffer (8.3% Dulbecco’s Modified Eagle Medium (DMEM), 24 mM NaHCO3, 2 mM glutamine) with 10 mM HEPES and 0.20% BSA and the islets were incubated at 37°C in for 1.5 hours. After incubation, the plate was centrifuged at RT, 1,000RPM for 1 minute. The 2.5 mM glucose-containing buffer was replaced with either, 2.5 mM, 9 mM, 16.7 mM or 22.4 mM glucose-containing buffer and the islets were incubated at 37°C for 1 hour. After incubation, 200uL of media was collected for secreted insulin quantification. Total insulin was extracted from the islets via free-thawing in 0.05% Triton. Both, media and extracts were stored at −20°C until downstream analysis. Secreted and total insulin was quantified using the Rat-High Range Insulin ELISA kit (ALPCO #80-INSRTH-E01, Salem, NH, USA), per manufacturer’s instructions. Secreted insulin is presented as % of content (total islet insulin).

### Glucose Tolerance Test

Glucose tolerance was assessed in male-only mice at P30. Mice were weaned at P21. Mice were fasted for 6 hours (∼8:00am – 2:00pm). 45 minutes prior to the glucose tolerance test (GTT), mice received a 200uL subcutaneous injection of 0.9% NaCl solution to prevent dehydration. Mice received an intraperitoneal injection of glucose at 2g/kg bodyweight. For blood glucose measurements, a blood sample was collected from the tail vein and quantified using the Bayer Contour® Blood Glucose Meter (Ascensia Diabetes Care #9545C, Basel, Switzerland) at time 0 (basal), 15, 30, 45, 60, 120 minutes post-D-glucose (Sigma-Aldrich #8270, St. Louis, MO, USA) intraperitoneal injection. For plasma insulin measurements, blood was collected from the tail vein using a Drummond™ Hemato-Clad™ Mylar™-Wrapped Hematocrit Tube (Drummond Scientific Company #1-00-7500-HC/5, Broomall, PA, USA), immediately, dispensed into a Heparin/Lithium Coated tube (Beckman Coulter #652825, Indianapolis, Indiana, USA) and placed on ice. To obtain plasma, blood was centrifuged at 11,800 x g for 1 min at RT. The plasma-containing supernatant was aliquoted and stored at −80°C until analysis. Insulin was quantified using the Mouse Ultrasensitive Insulin ELISA (ALPCO #80-INSMSU-E01, Salem, NH, USA), per manufacturer’s instructions.

### RNA isolation and RT-qPCR

Total RNA was isolated from ∼400 islets using the RNeasy Plus Micro Kit (QIAGEN #74034, Venlo, Netherlands) per the manufacturer’s instructions. Each total RNA sample was quantified using a spectrophotometer (Thermo Scientific Nanodrop™ 1000, Waltham, MA, USA). 500 ng of total RNA was reverse transcribed using the iScript cDNA Synthesis Kit (Bio-Rad #1708891, Hercules, CA, USA) per the manufacturer’s instructions. 10 ng of cDNA was used to estimate the abundance of specific mRNA transcripts using the iTaq Universal SYBR Green Supermix (Bio-Rad #1725121, Hercules, CA, USA) on a CFX96 Touch Real-Time PCR Detection System (Bio-Rad, CA). Relative mRNA transcript abundance was estimated using the ΔΔCt method [23] with the housekeeping genes, *18S* and *Rplp0*, used as an internal normalization control for each sample. Statistical analysis was performed on delta CT data. Primer sequences used are provided in **Supplementary Table 1**.

### Dapagliflozin Treatment

To study the effect of hyperglycemia on the phenotypic progression, we utilized dapagliflozin intervention to reduce hyperglycemia. This intervention was initiated on P21, the day of weaning. Mice were provided with Dapagliflozin-containing (Thermo Scientific, #AC464620010, Waltham, MA, USA), water as an exclusive water source, from P21 until euthanasia at P40 and maintained on standard chow. Dapagliflozin dose was 5mg/kg/day and assessed by daily water consumption of 4mL/25g bodyweight [24]. The treatment continued until P40, at which fasting blood glucose, fasting plasma insulin, and histological analysis were conducted, as described above.

### Islet Glutathione Measurement

Simultaneous quantification of islet GSH and GSSG was conducted by liquid chromatography mass spectrometry (LC-MS/MS), as previously described [25]. Islets were isolated, as described above. ∼100 isolated islets were centrifuged at 4°C, 1,500 RPM for 3 minutes, supernatant removed, and resuspended in 500uL of *N*-Ethylmaleimide (NEM) (Millipore Sigma #E3876, Burlington, MA, USA) solution (2.5 mM NEM in 80% Methanol) for simultaneous thiol alkylation of free GSH and protein precipitation for 20 minutes at RT. Incubated islets were then, centrifuged at 4°C for 10 minutes, 13,000g. The supernatant was dried, overnight, in a SpeedVac™ Concentrator (Thermo Scientific #SPD111V, Waltham, MA, USA) and resuspended in 100uL 50 mM ammonium acetate, to which, 300uL of dichloromethane (MilliporeSigma Supleco, Burlington, MA, USA) was added to extract excess NEM via 1 minute vortex and centrifugation at 4°C for 5 minutes, 13,000g. The protein pellet was flash frozen for downstream quantification by BCA (described above). The upper phase was aliquoted, diluted 1 to 5 in 50 mM ammonium acetate, and 2uL was injected for LC-MS/MS detection. LC-MS/MS detection was conducted on a UPLC Acquity I-Class (Waters Corporation, Milford, USA) interfaced with a Xevo TQ-S micro triple quadrupole mass spectrometer (Xevo TQ-S-micro, Waters Corporation, Milford, USA) using the following LC conditions: Acquity UPLC HSS T3 1.8uM 2.1 x 100mm (Waters Corporation #186003539, Milford, USA) column with solvent (solvent A 0.1% formic acid in water and solvent B 0.1% formic acid in acetonitrile) gradient (0-10 min 1-30% B, 10-15 min 30-70% B, and 15-20min 1% B) at a 0.3mL/min flow rate. Mass spectrometry detection occurred in positive-mode using multiple reaction monitoring (MRM) transitions for derivatized GSH. Multiple reaction monitoring (MRM) transitions are reported in **Supplementary Table 2**. MS was equipped with electrospray ionization (ESI) source and detector was used in positive mode. The MS was operated under the following parameters: capillary voltage 3kV, source temperature 150°C, desolvation gas temperature 350°C, cone gas flow 50 L/hr, desolvation gas flow 400 L/hr. Additionally, a standard curve for derivatized GSH was prepared and analyzed, in-parallel to support quantification of GSH (see **Standard Curve Solution Preparation**). All solvents used were LC-MS grade (Fisher Chemical, Hampton, NH, USA), unless otherwise specified. Data was acquired using MassLynx version 4.1 (Waters Corp., Milford, MA, USA). TargetLynx (Waters Corp., Milford, MA, USA) was used to integrate mass spectrum of identified compounds. Data was normalized to protein concentration.

### Standard Curve Solution Preparation

A stock solution of 10 mM GSH was prepared in water and immediately, directly diluted in 2.5 mM NEM in 50 mM ammonium acetate to 100 uM GSH. The 100 uM GSH-NEM solution was incubated for 20 minutes at RT to allow for thiol alkylation of free GSH. 100 uM GSH-NEM was serially diluted in 50 mM ammonium acetate to generate a standard curve at 25.00 (7.500), 10.00 (3.000), 5.000 (1.500), 2.500 (0.750), 1.250 (0.375), 0.625 (0.187), 0.313 (0.094), 0.156 (0.047), 0.078 (0.023), and 0.039 (0.011) uM (ug/mL). The standard curve was analyzed in triplicate, prior to analyzing sample.

### Statistical Analysis

Data are presented as mean ± standard deviation. A two-tailed, unpaired, *t* test was used to perform comparisons between two groups. A one-way ANOVA and Tukey’s post hoc test was used to perform comparisons between three or more groups. Significance determined at *P* value < 0.05. Data were analyzed using GraphPad Prism v9.0 (GraphPad, San Diego, CA, USA).

## Results

### Gclc^f/f^/Le-Cre^Tg/-^ mice (Gclc KO) display an age-related, progressive diabetes phenotype

Preliminary health assessment indicated that *Gclc* KO mice died or required euthanasia (due to poor health) at approximately 1.5 months (∼45 - 50 days) of age. Prior to death, *Gclc* KO mice presented with increased urination (indicated by wet bedding and a pungent smell), hunched posture, and visible weight loss. These symptoms were similar to uncontrolled diabetes and suggested that *Gclc* KO mice might be diabetic. Indeed, fasting blood glucose levels in *Gclc* KO mice aged P40 (∼524 mg/dL) were significantly higher, when compared to *Gclc* CTRL (∼163 mg/dL) **(1b)**. Because we sought to determine the cause not effect of fasting hyperglycemia in our model, we focused our investigations on *Gclc* KO mice aged P40 and younger. Diabetes is a progressive condition therefore, we assessed fasting blood glucose levels as a function of age in *Gclc* KO mice. We expanded our initial assessment to include the pre-weaning (P14), newly weaned (P21), early post-weaning (P25), and mid to late post-weaning (P30 and P35) periods. *Gclc* KO mice displayed age-related, progressive fasting hyperglycemia **(1b)**. Over the pre-weaning (P14), newly weaned (P21) and early post-weaning (P25) period, fasting blood glucose levels did not significantly differ, when compared to *Gclc* CTRL. Over the mid to late post-weaning period (P30, P35, and P40), *Gclc* KO mice presented with hyperglycemia and fasting blood glucose was significantly higher, when compared to *Gclc* CTRL. The severity of the fasting hyperglycemia increased with age, from moderate at P30 (∼250 mg/dL vs. Gclc CTRL ∼178 mg/dL) to severe at P35 (∼460 mg/dL vs. Gclc CTRL ∼189 mg/dL) thru P40 (∼526 mg/dL vs. Gclc CTRL ∼169 mg/dL). The activation of the Pax6 gene promoter in pancreatic endocrine progenitor cells at ∼E9.5 facilitated the serendipitous deletion of *Gclc* gene expression (confirmed by protein expression and gene expression in primary islets isolated from P21 *Gclc* KO mice **(6a and 6c)**). Insulin resistance is associated with fasting hyperglycemia. However, the islet-specific deletion of *Gclc* hinted at impaired islet-function, namely β cell insulin secretion, and consequential reduction in plasma insulin as causal for fasting hyperglycemia. Therefore, we assessed fasting plasma insulin levels as a function of age in *Gclc* KO mice. Hypoinsulinemia, evidenced by significantly lower fasting plasma insulin levels in *Gclc* KO mice when compared to *Gclc* CTRL, was observed in severely hyperglycemic (P35 and P40) *Gclc* KO mice **(1c)**. Interestingly, fasting plasma insulin levels in moderately hyperglycemia *Gclc* KO mice (P30) did not significantly differ from *Gclc* CTRL. The hyperglycemia and hypoinsulinemia are independent of a significant difference in the bodyweight of *Gclc* KO when compared to *Gclc* CTRL mice **(1a)**.

**Fig. 1a-c.**
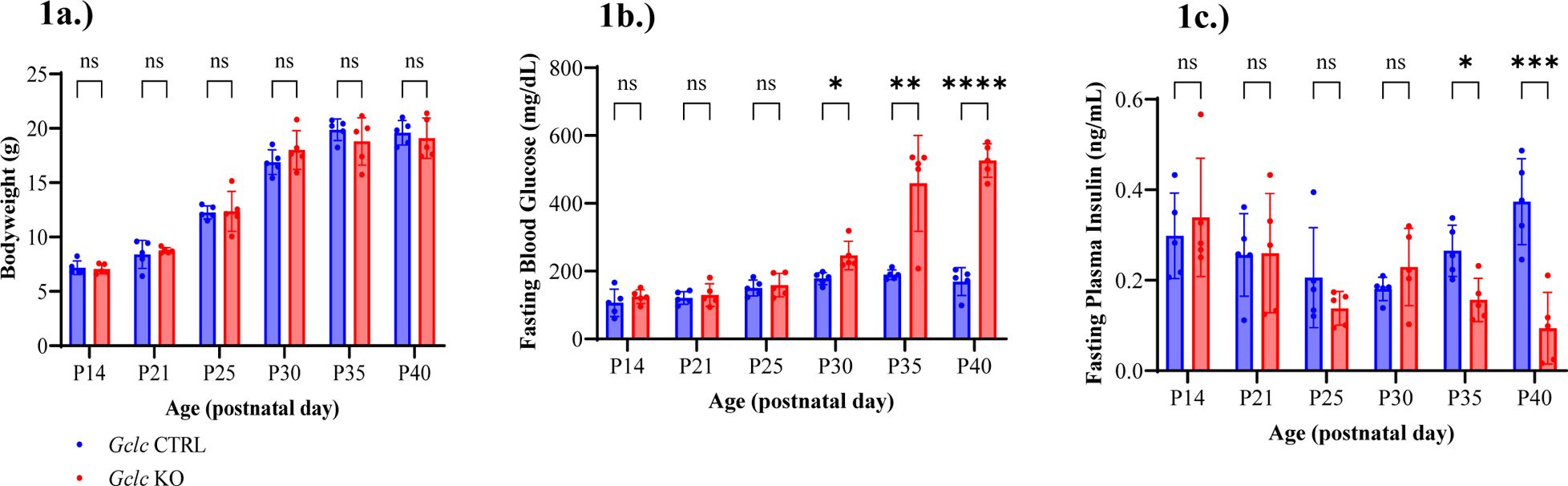
Gclc^f/f^/Le-Cre^Tg/-^ mice (Gclc KO) display an age-related, progressive diabetes phenotype. Physiological characterization of *Gclc* KO compared to *Gclc^f/f^*/*Le-Cre^−/−^* (*Gclc* CTRL) mice at postnatal day (P) 14, 21, 25, 30, 35, and 40. **a.)** Bodyweight (g) **b.)** 6-hr morning fast (∼8:00am – 2:00pm), fasting blood glucose (mg/dL) **c.)** 6-hr morning fast (∼8:00am – 2:00pm), fasting plasma insulin (ng/mL). Results are presented as mean ± S.D., n = 5 mice/genotype. Statistical analysis was performed using an unpaired t-test. Significance was determined at *p-value < 0.05, **, ***, ****p-value < 0.01, ns = non-significant.

### Moderately hyperglycemic Gclc KO mice display impaired glucose homeostasis

Phenotypic characterization of *Gclc* KO mice was improved by performing glucose tolerance tests (GTT) to assess glucose homeostasis. At P30, *Gclc* KO mice were moderately hyperglycemic and did not display significantly lower fasting plasma insulin therefore, we performed our GTTs at P30 to determine if impaired glucose homeostasis was coincident with impaired β cell function. Moderately hyperglycemic *Gclc* KO mice were glucose intolerant but, plasma insulin levels did not significantly differ after glucose challenge. *Gclc* KO mice presented with significantly higher blood glucose levels at time 0, 15, 30, 45, 60, and 120 minutes, and area under the curve (AUC) for blood glucose, compared to *Gclc* CTRL **(2a-b)**. Additionally, *Gclc* KO mice presented with a non-significant decrease in AUC for plasma insulin, compared to *Gclc* CTRL **(2c-d)**.

**Fig. 2a-d.**
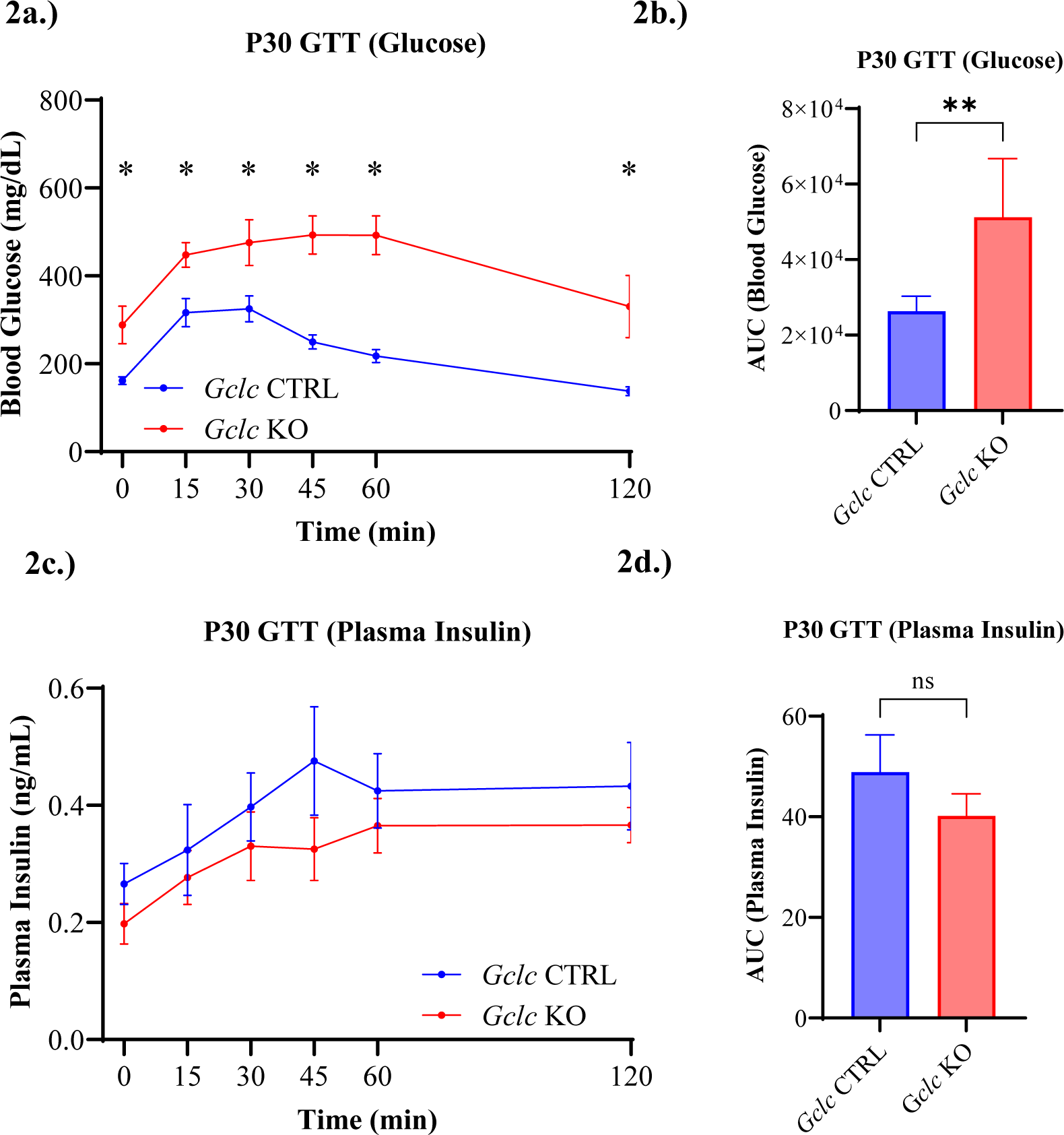
Moderately hyperglycemic Gclc *KO* mice display impaired glucose homeostasis. Glucose tolerance testing (GTT), after a 6-hr morning fast (∼8:00am – 2:00pm), was used to determine glucose homeostasis in *Gclc* KO mice at postnatal day (P) 30, prior to severe hyperglycemia. **a-b.)** P30 GTT blood glucose **(a)**, area under curve (AUC) blood glucose **(b) c-d.)** P30 GTT plasma insulin **(c)**, AUC plasma insulin **(d)**. Results are presented as mean ± S.E.M., n = 5 mice/genotype. Statistical analysis was performed using an unpaired t-test on AUC ± S.E.M., or mean ± S.D for comparison at individual timepoints. Significance was determined at *p-value < 0.05, ****p-value < 0.01, ns or no * = non-significant.

### Progressive histopathological changes are present in the pancreatic islets of Gclc KO mice

While plasma insulin levels in response to glucose challenge did not significantly differ, given the islet-specific *Gclc* deletion and the impairment to glucose homeostasis in moderately hyperglycemic mice, we hypothesized that islet-specific pathology might predate severe hyperglycemia. We performed qualitative histological and immunohistochemical assessment of *Gclc* KO mice at P21, P25, P30, P35, P40 **(3a-c)**. Over the newly weaned (P21) and early post-weaning (P25) period, *Gclc* KO islets were enlarged and non-circular but, displayed uniform insulin and glucagon expression. Over the mid to late post-weaning period (P30, P35, and P40), *Gclc* KO islets displayed progressive vacuolization and reduced islet mass with the progressive loss of insulin and glucagon expression.

**Fig. 3a-c.**
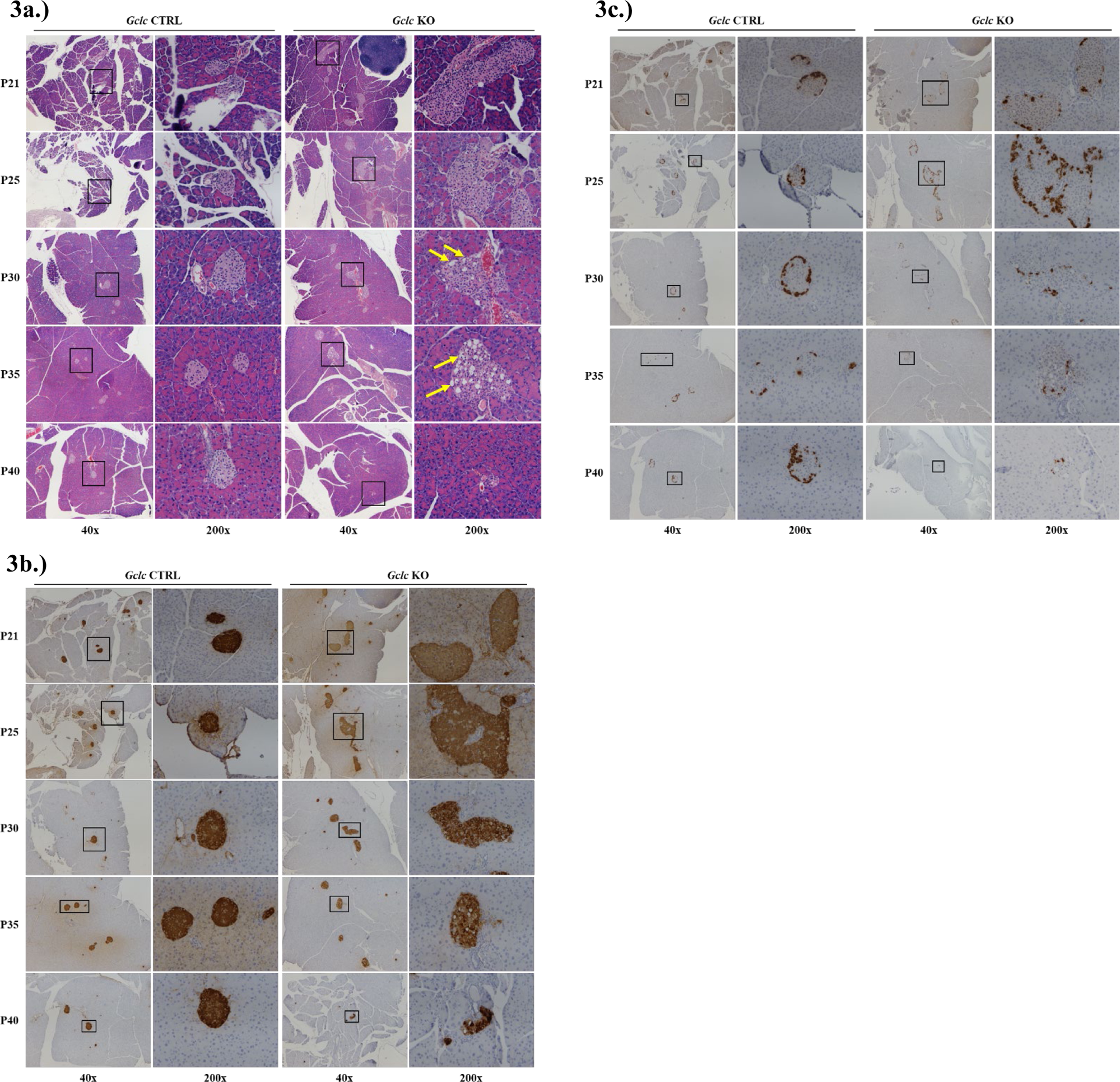
Progressive histopathological changes are present in the pancreatic islets of Gclc KO mice. Representative images of stained pancreata from *Gclc* CTRL and *Gclc* KO mice at postnatal day (P) 21, 25, 30, 35, 40. **a.)** Hematoxylin & Eosin staining **b.)** Immunohistochemical staining for Insulin **c.)** Immunohistochemical staining for Glucagon. Islet-regions indicated by black box. Vacuolation indicated by yellow arrows.

### Lowering blood glucose does not prevent the diabetes phenotype in Gclc KO mice

Glucotoxicity, due to chronic hyperglycemia, is associated with islet-specific vacuolization and eventual, cell death. We observed islet-specific vacuolization, alongside hyperglycemia. Thus, we hypothesized that glucotoxicity was causal for islet-specific histopathological changes and diabetes phenotype. To test this hypothesis, we treated *Gclc* KO and *Gclc* CTRL mice with Dapagliflozin, an SGLT2 inhibitor, to promote the renal loss of glucose and hence lower blood glucose levels. Mice were administered Dapagliflozin via drinking water from weaning at P21 until euthanasia at P40. Dapagliflozin treatment was successful and fasting blood glucose levels were significantly lower in treated *Gclc* KO mice when compared to untreated *Gclc* KO mice **(4a)**. Dapagliflozin treatment did not significantly lower fasting blood glucose levels in treated *Gclc* CTRL when compared to untreated *Gclc* CTRL (p = 0.746). While Dapagliflozin treatment successfully reduced fasting blood glucose levels, treated *Gclc* KO mice remained hypoinsulinemic **(4b)** and fasting plasma insulin levels did not significantly differ when compared to untreated *Gclc* KO mice suggesting that a defect in insulin synthesis was inherent to the *Gclc* KO β cell and not dependent upon a respond to the blood glucose level. Dapagliflozin treatment did not significantly lower fasting plasma insulin levels in treated *Gclc* CTRL when compared to untreated *Gclc* CTRL. Clinically, Dapagliflozin treatment is associated with weight loss however, treatment of mice with Dapagliflozin did not significantly decrease bodyweight **(4c)**. We examined pancreata of treated *Gclc* KO mice for rescue of islet-specific vacuolization. Dapagliflozin treatment did not prevent islet-specific vacuolization **(4d)**. The inability of Dapagliflozin treatment to prevent hypoinsulinemia and islet-specific vacuolization suggested that, glucotoxicity is not the primary driver of the diabetes phenotype.

**Fig. 4a-d.**
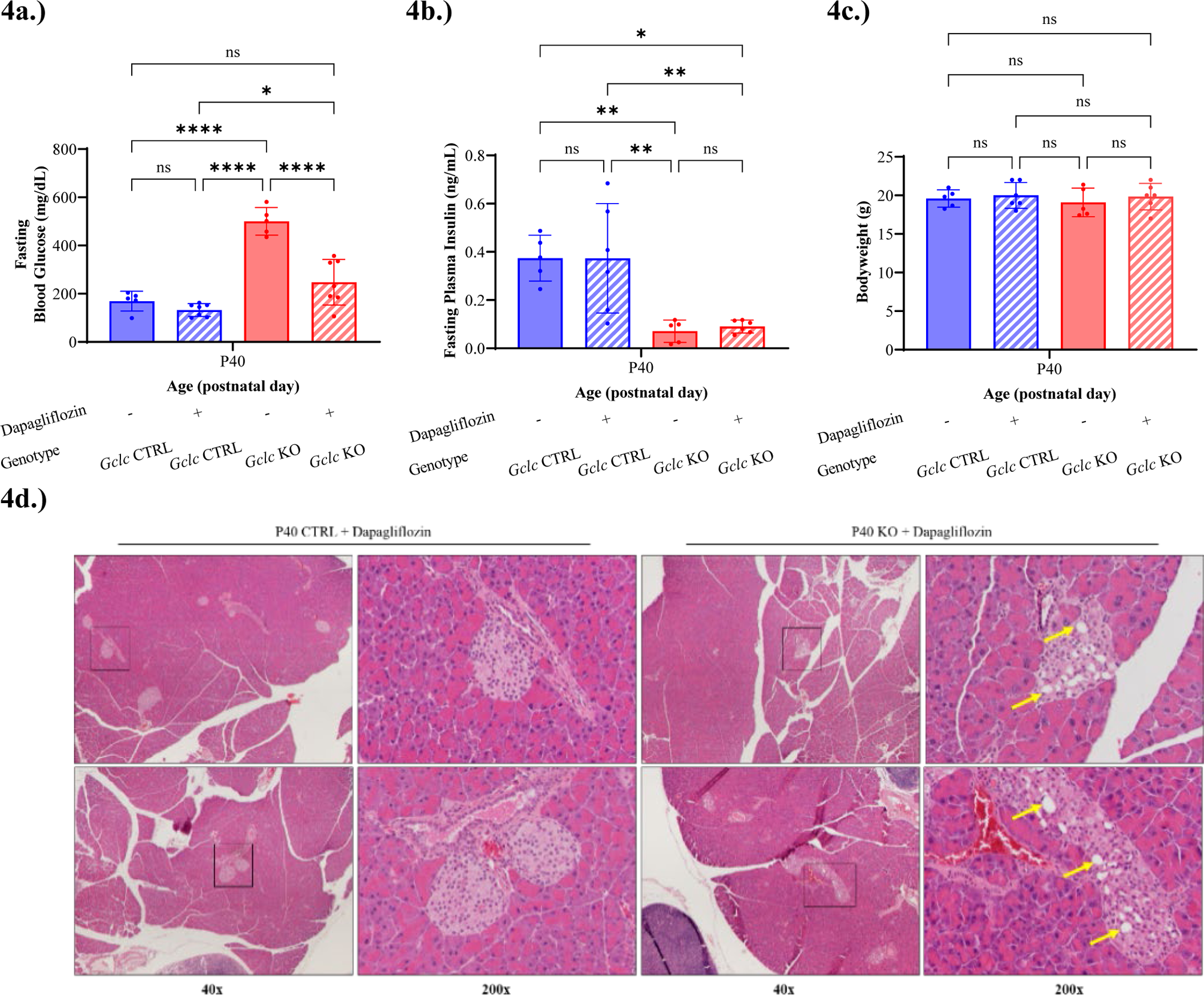
Lowering blood glucose does not prevent the diabetes phenotype in Gclc KO mice. Mice were provided Dapagliflozin treated drinking water from postnatal day (P) 21 to P40, as described in *Materials and Methods*. At P40, differences in the fasting blood glucose, fasting plasma insulin, and bodyweight of untreated (*Gclc* CTRL and *Gclc* KO) and treated (*Gclc* CTRL + Dapagliflozin and *Gclc* KO + Dapagliflozin) were compared. **a.)** 6-hr morning fast (∼8:00am – 2:00pm), fasting blood glucose (mg/dL) **b.)** 6-hr morning fast (∼8:00am – 2:00pm), fasting plasma insulin (ng/mL) **c.)** Bodyweight (g) **d.)** Representative images of Hematoxylin & Eosin stained pancreata from P40 *Gclc* CTRL + Dapagliflozin and P40 *Gclc* KO + Dapagliflozin mice. Islet-regions indicated by black box. Vacuolation indicated by yellow arrows. Results are presented as mean ± S.D., n = 5-6 mice/genotype/treatment group. Statistical analysis was performed using a Two-Way ANOVA with Tukey’s Post-Hoc test. Significance was determined at *p-value < 0.05, **, ****p-value < 0.01, ns or no = is non-significant.

### Newly weaned Gclc KO mouse islets are enlarged but, display impaired glucose stimulated insulin secretion

Qualitative histopathological assessment revealed newly weaned (P21) *Gclc* KO mouse islets were enlarged when compared to the *Gclc* CTRL mouse islets. We quantitatively confirmed this observation and, on average, *Gclc* KO mouse islets are significantly larger than *Gclc* CTRL mouse islets **(5b)**. The significant increase in *Gclc* KO islet size does not coincide with a significant increase in total islet mass or number **(5a,c)**. The neonatal period, at weaning and after, requires an increase in islet size to support an increase in insulin demand during rapid postnatal growth. We assessed the functionality of newly weaned islets, *ex vivo*, by performing GSIS assays on primary isolated islets. As expected, *Gclc* CTRL mouse islets exhibited a dose-response increase in insulin secretion when challenged with an increasing glucose load. Surprisingly, *Gclc* KO mouse islets, although enlarged, were dysfunctional **(5d)**. Insulin secretion did not significantly differ between *Gclc* KO and *Gclc* CTRL mouse islets when exposed to 2.5 mM (basal) or 9 mM (stimulatory) glucose load. However, when challenge with 16.7 mM and 22.4 mM glucose load, *Gclc* KO mouse islets secreted significantly less insulin when compared to *Gclc* CTRL. Functional changes were independent of significant differences in total islet insulin content between *Gclc* KO and *Gclc* CTRL mice **(5e)**. We sought to determine the mechanistic cause for impaired glucose stimulated insulin secretion and characterized the expression of genes required for islet cell identity and insulin secretion. These included: β cell identity (*Ins1*, *Ins2*, *Pdx1*, *Nkx6.1*, *MafA*, *NeuroD*), α cell identity (*Gcg*, *MafB*), δ cell identity (*Sst*), whole islet identity (*Pax6*, *Ngn3*), islet ductal identity (*Hnf6*), and glucose stimulated insulin secretion (*Glut2*, *Gck*, *Abcc8*, *Kcnj11*, *Cacnald*, *Fox01*) **(5f)**. The expression of β cell, α cell, and δ cell hormone genes were significantly downregulated. However, the expression of cell identity-related transcription factors, which regulate hormone gene expression, were not significantly downregulated. Additionally, the expression of islet ducal identity and glucose stimulated insulin secretion genes were unchanged.

**Fig. 5a-f.**
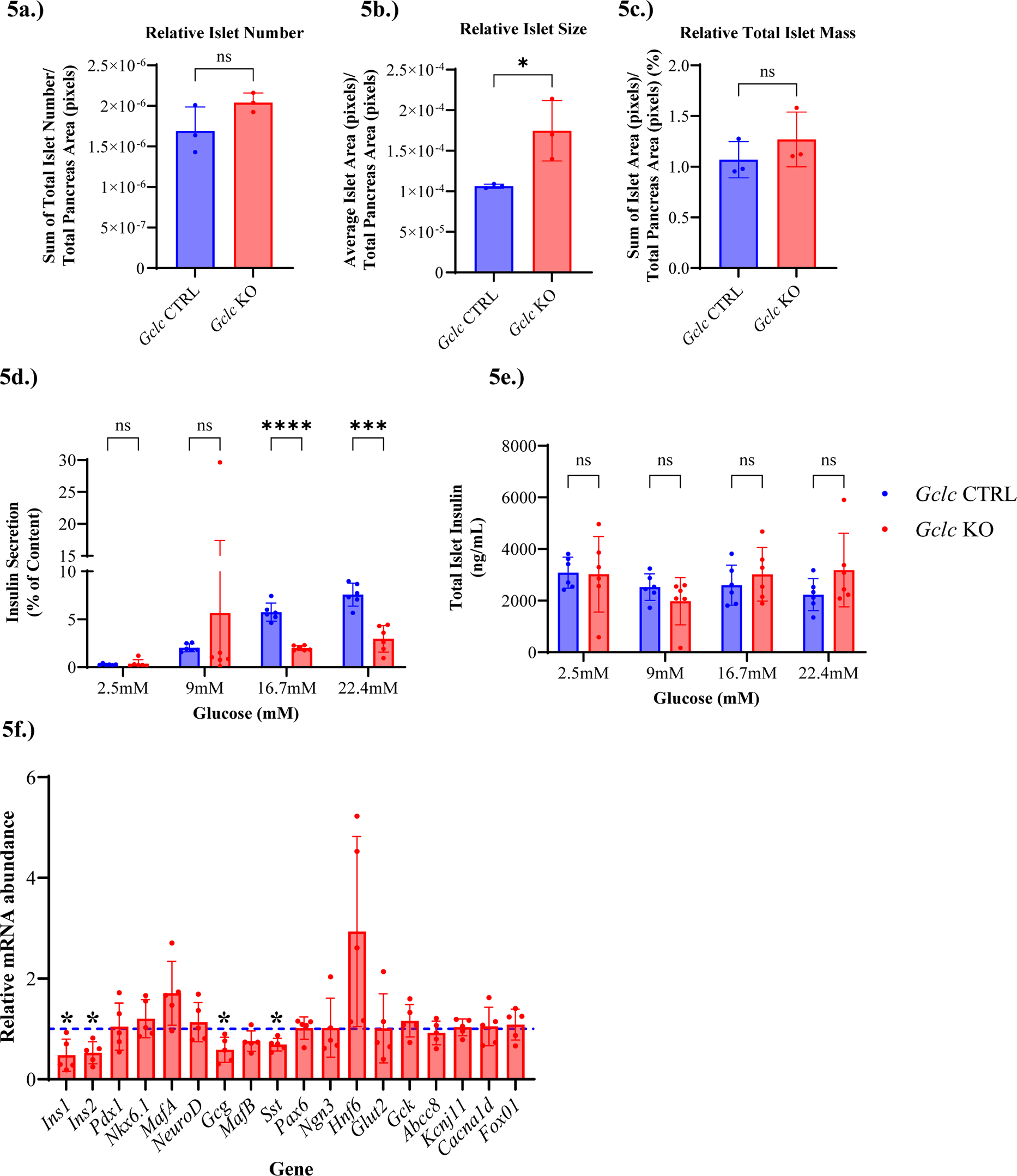
Newly weaned Gclc KO mouse islets are enlarged but, display impaired glucose stimulated insulin secretion. **a.)** relative islet number in pancreas at postnatal day (P) 21 **b.)** relative islet size in pancreas at P21 **c.)** relative total islet mass in pancreas at P21. Islets (64-124 islets/mouse) were quantified by Hematoxylin & Eosin staining (n = 3 mice/genotype). **d.)** glucose-stimulated insulin secretion (% of content) in isolated islets from P21 mice **e.)** total islet insulin content (ng/mL) in isolated islets from P21. Each replicate (n = 6 replicates/genotype/glucose condition), represents 10-islets, seeded in one-well of a 96-transwell plate, derived from a pool of ∼400 islets, isolated from the pancreata of 3-4 mice/genotype. **f).** qPCR analysis of relative gene expression in isolated islets from P21 mice. Housekeeping genes *18s* and *Rplp0* were used for normalization of cycle-threshold (CT) data. The relative mRNA abundance of an individual gene is reported as fold of the *Gclc* CTRL. Each replicate (n = 5 replicates/genotype) represents a pool of ∼400 islets, isolated from the pancreata of 3-4 mice/genotype. All results are presented as mean ± S.D. Statistical analysis was performed using an unpaired t-test. For q-PCR analysis, statistical analysis was performed on delta CT data. Significance was determined at *p-value < 0.05, ***, ****p-value < 0.01, ns or no = is non-significant.

### Newly weaned KO mouse islets have diminished islet GSH biosynthesis and are undergoing oxidative stress

We confirmed the deletion of *Gclc* by quantifying *Gclc* gene expression and GCLC protein expression in primary isolated islets from newly-weaned (P21) mice. *Gclc* gene **(6c)** and GCLC protein expression **(6a)** were significantly decreased in *Gclc* KO mouse islets, when compared to *Gclc* CTRL mouse islets. Diminished islet GSH biosynthesis is evidenced by the significantly lower reduced GSH concentration in *Gclc* KO mouse islets, when compared to *Gclc* CTRL mouse islets **(6b)**. The depletion of reduced GSH can induce oxidative stress. Therefore, we sought to determine if newly weaned *Gclc* KO mouse islets were undergoing oxidative stress and measured the expression of oxidative stress-response genes. These include genes of the two major cellular redox systems, GSH (*Gclm, Gsr, Gpx1, and Gpx4*) and thioredoxin (Trx) (*Txn1, Txn2, Txnrd1, and Txnip*) and antioxidant defense (*Nrf2, Hmox1*, *Nqo1*, *Bach1*) **(6c)**. The deletion of *Gclc* did not confer increased gene expression of *Gclm*, *Gpx1*, or *Gsr*. However, *Gpx4* gene expression was significantly upregulated in newly weaned *Gclc* KO mouse islets. Of the antioxidant defense genes, *Nrf2* gene expression was not upregulated but, the expression of *Nrf2*-regulated genes, *Hmox1* and *Nqo1*, were significantly upregulated in newly weaned *Gclc* KO mouse islets. This upregulation suggests, increased nuclear translocation and activity of NRF2 via the NRF2-antioxidant response element pathway. While the thioredoxin system is, typically, a back-up for the GSH system to support cellular redox state, the expression of Trx system genes were not significantly upregulated in newly weaned *Gclc* KO mouse islets.

**Fig. 6a-b.**
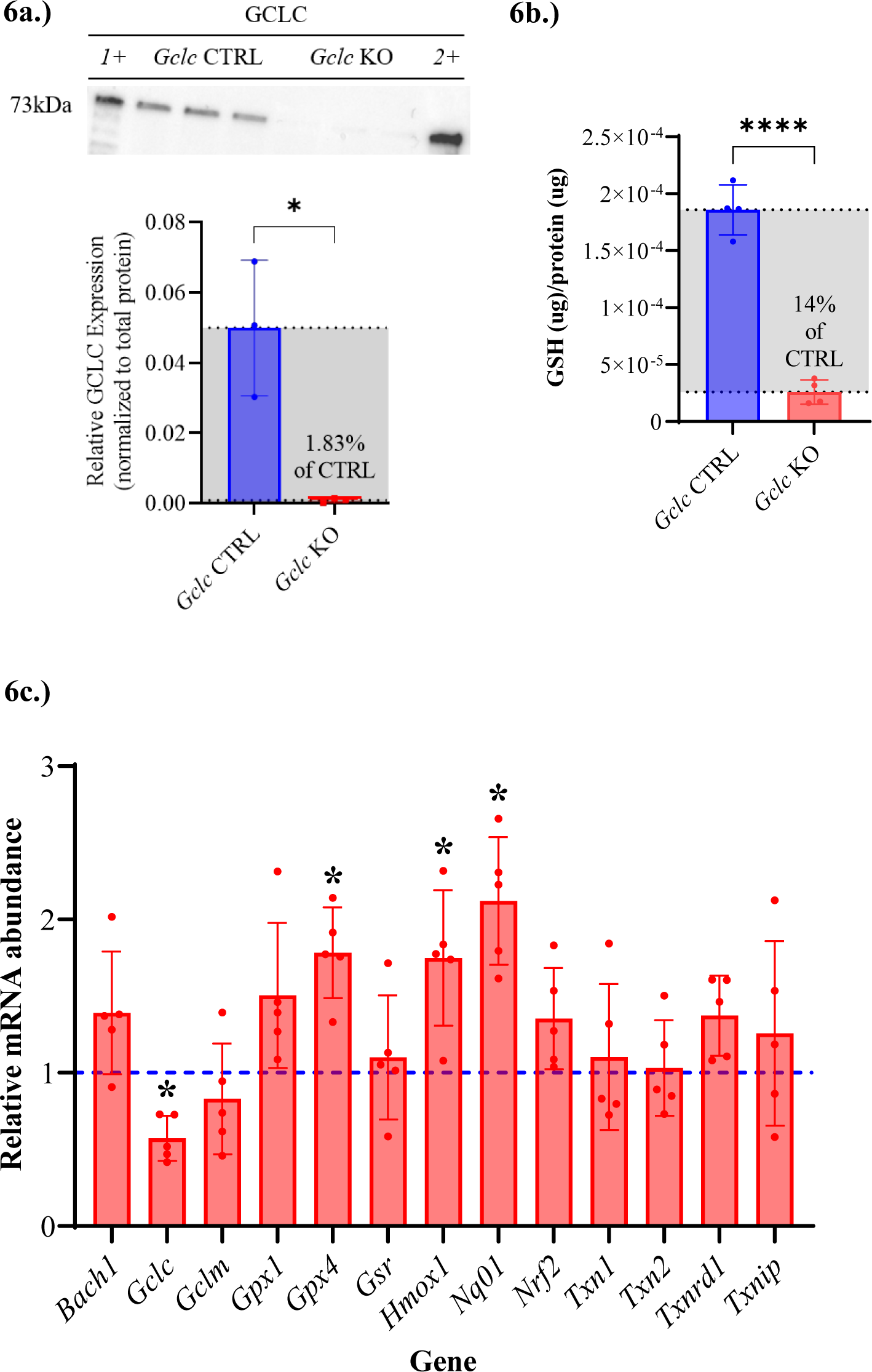
Newly weaned Gclc KO mice have diminished islet GSH biosynthesis and are undergoing oxidative stress. **a.)** Western blotting image for GCLC expression in isolated islets from postnatal day (P) 21 mice. Protein levels were quantified by densitometry analysis and normalized to total protein loading per sample. 1+ and 2+ indicate positive controls from liver for GCLC. Each replicate (n = 3 replicates/genotype) represents a pool of ∼400 islets, isolated from the pancreata of 3-4 mice/genotype. **b.)** the concentrations of reduced (GSH) was measured in isolated islets from P21 mice using targeted liquid chromatography mass spectrometry and normalized to total protein (ug). Each replicate (n = 4 replicates/genotype) represents a pool of ∼100 islets, isolated from the pancreata of 3-4 mice/genotype. **c.)** qPCR analysis of relative gene expression in isolated islets from P21 mice. Housekeeping genes *18s* and *Rplp0* were used for normalization of cycle-threshold (CT) data. The relative mRNA abundance of an individual gene is reported as fold of the *Gclc* CTRL. Each replicate (n = 5 replicates/genotype) represents a pool of ∼400 islets, isolated from the pancreata of 3-4 mice/genotype. All results are presented as mean ± S.D. For q-PCR analysis, statistical analysis was performed on delta CT data. Statistical analysis was performed using an unpaired t-test. Significance was determined at *p-value < 0.05, ***, ****p-value < 0.01, ns or no * = non-significant.

### Newly weaned Gclc *KO* mouse islets display characteristics of premature senescence

Examination of newly weaned *Gclc* KO mouse islets revealed significantly enlarged yet, dysfunctional islets with significantly decreased islet hormone gene expression. The significant enlargement of islet size can be driven by hyperplasia (increased cell number) or hypertrophy (increased cell size). We characterized the expression of cell cycle-related genes including: cell cycle inhibitors, p53 (*Trp53)*, p21 (*Cdkn1a*) and p16 (*Ckdn2a*), and the marker of proliferation, *Mki67* **(7a)**. While the expression *Trp53* and *Ckdn2a* were not significantly changed, the expression of *Cdkn1a* was significantly upregulated in newly weaned *Gclc* KO mouse islets, when compared to *Gclc* CTRL. Surprisingly, the expression of *Mki67* was unchanged, when compared to *Gclc* CTRL. Given these unexpected results, we re-examined the histopathology of newly weaned *Gclc* KO mouse islets and discovered hypertrophic islet regions **(7b)**. Together, cell cycle arrest and cellular swelling (hypertrophy) are characteristic of cellular senescence [26]. Therefore, we examined the expression of genes, related to the Senescence-Associated Secretory Phenotype (SASP) and pro-survival Senescence Cell Anti-Apoptotic Pathways (SCAPs). These genes included: cytokines (*Il-6* and *Tnf-α*), extracellular proteases (*Mmp3* and *Timp1*), and pro-survival (*Bcl2*) **(7a)**. Of those genes, the expression of Il-6 was significantly higher in newly weaned *Gclc* KO islets, when compared to *Gclc* CTRL.

**Fig.7a-b.**
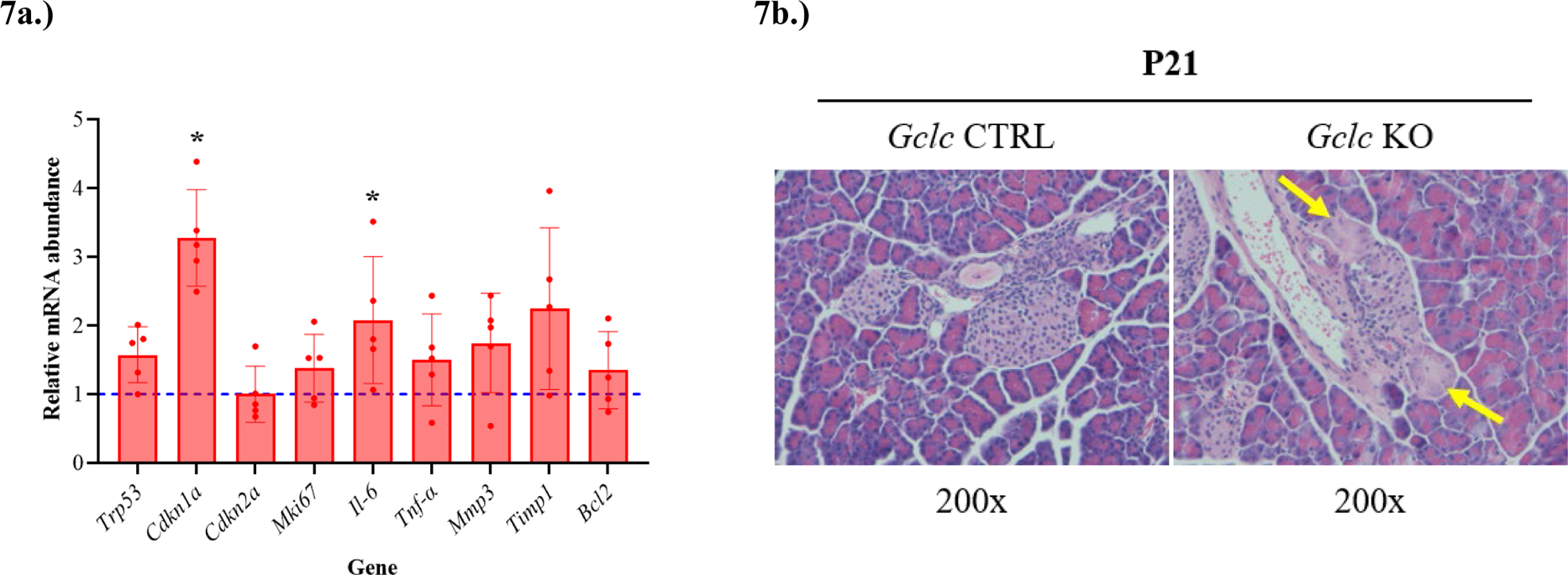
Newly weaned Gclc KO mice display characteristics of premature senescence. **a.)** qPCR analysis of relative gene expression in isolated islets from postnatal day (P) 21 mice. Housekeeping genes *18s* and *Rplp0* were used for normalization of cycle-threshold (CT) data. The relative mRNA abundance of an individual gene is reported as fold of the *Gclc* CTRL. Each replicate (n = 5 replicates/genotype) represents a pool of ∼400 islets, isolated from the pancreata of 3-4 mice/genotype. **b.)** Representative images of Hematoxylin & Eosin stained pancreata from *Gclc* CTRL and *Gclc* KO mice at P21. Hypertrophic islet-regions indicated by yellow arrows. Results are presented as mean ± S.D. Statistical analysis was performed on delta CT data. Statistical analysis was performed using an unpaired t-test. Significance was determined at *p-value < 0.05, ***, ****p-value < 0.01, ns or no * = non-significant.

## Discussion

We provide the first characterization of a diabetes phenotype observed in *Gclc* KO mice wherein, the deletion of the *Gclc* gene is facilitated by *Pax6* promoter P0 mediated Cre expression during pancreatic development. Our results demonstrate that, *Gclc* KO mice display an age-related, progressive diabetes phenotype. Prior to severe hyperglycemia at P35 and P40, *Gclc* KO mice with moderate hyperglycemia are glucose intolerant. Histological examination of pancreata revealed age-related islet-specific pathology in *Gclc* KO mice including: enlarged islets in the newly weaned (P21) period, followed by worsening islet-specific cellular vacuolization, decreasing islet hormone protein expression, and progressive loss of islet mass over the mid to late post-weaning (P30, P35, P40) period. The inability of lowered blood glucose to rescue the diabetes phenotype suggested intrinsic islet defects, predate the diabetes phenotype. Therefore, we functionally and molecularly characterized the enlarged islets of newly weaned (P21) *Gclc* KO mice. Islets from newly weaned (P21) *Gclc* KO mice had impaired GSIS, reduced islet hormone expression, were deficient in GSH, and undergoing oxidative stress. Investigations to determine the cause for islet enlargement revealed significant cell cycle inhibition and increased markers of senescence. Collectively, this data demonstrates the essentiality of GSH biosynthesis in endocrine pancreas development and suggest that this essentiality is related to developmental cell fate.

We observed chronic hyperglycemia, increasing in severity, in *Gclc* KO mice of mid to late post weaning age (P30, P35, P40). Chronic hyperglycemia can elicit glucotoxicity. Glucotoxicity is defined by impaired glucose tolerance with decreased insulin secretion, the eventual loss of insulin content and in turn, blood insulin [27]. We observed impaired glucose tolerance at P30 followed by, the progressive decrease in fasting plasma insulin (P40), in *Gclc* KO mice. In addition to eliciting β cell dysfunction, glucotoxicity can cause islet cell death [28]. Vacuolization, as observed in *Gclc* KO mice at P30, P35, and P40, is a morphology, which is associated with various forms of cell death [29]. Our results suggested that, *Gclc* KO mouse islets are undergoing cell death possibly, via glucotoxicity. We employed an SGLT2 inhibitor to significantly reduce fasting blood glucose and thus, determine if glucotoxicity was causal for the diabetes phenotype. Reduction in fasting blood glucose did not rescue the diabetes phenotype, as evidenced by the decrease in fasting plasma insulin and islet-specific vacuolization when compared to *Gclc* CTRL. This demonstrates that the cytotoxic stimulus for beta cell dysfunction and cellular vacuolization is not driven by hyperglycemia. In deleting *Gclc*, we significantly reduced GSH biosynthesis. GSH depletion can induce a variety of cell death processes including but, not limited to apoptosis and necrosis [30, 31]. Based-on our data, we cannot fully determine the type of cell death process occurring in mid to late weaning *Gclc* KO mice. Improved characterized of the vacuolization would illuminate the cell death process.

The most significant nutritional challenge faced by a developing mammal is weaning. From birth, rodents must physiologically adapt to leverage the dietary transition from mother’s milk, a food matrix high in lipid and protein, to standard chow, a food matrix high in carbohydrate, for the rapid post-weaning growth period [32]. We observed that *Gclc* KO mice do not display overt diabetes until after weaning. This suggested that *Gclc* KO mice harbored dysfunctional islets and the manifestation of this dysfunctionality was elicited by increased insulin demand during post-weaning adaptation. Indeed, islets from newly weaned *Gclc* KO mice can be stimulated to secrete insulin, similarly to *Gclc* CTRL, with 9 mM glucose. However, when stimulated with 16.7 mM and 22.4 mM glucose load, *Gclc* KO islets cannot significantly increase insulin secretion. To investigate the mechanism for the diminished secretory response, we examined the expression levels of islet genes related to hormone expression (*Ins1*, *Ins2*, *Gcg*, *Sst*), β cell transcription factors (*Pdx1*, *Nkx6.1*, *MafA*, *NeuroD*), α cell transcription factors (*MafB*), general islet transcription factors (*Pax6*, *Ngn2*, *Hnf6*), glucose sensing and insulin secretion (*Glut2*, *Gck*, *Abcc8*, *Kcnj11*, *Cacnald*, *Fox01*). Surprisingly, the expression levels of most genes did not significantly differ between *Gclc* KO and *Gclc* CTRL. The islet cell hormones genes, *Ins1*, *Ins2*, *Gcg*, *Sst*, were all expressed at significantly lower levels in *Gclc* KO when compared to *Gclc* CTRL. The significant decrease in *Ins1* and *Ins2* gene expression cannot account for the diminished secretory response. Total insulin measured in *Gclc* KO islet after GSIS was not significantly different from *Gclc* CTRL. The synthesis and degradation of mRNA take place on a timescale more rapid than the synthesis and degradation of protein [33]. Therefore, if the decrease in *Ins1* and *Ins2* gene expression is maintained, the insulin content will eventually decrease. This is reflected in the *Gclc* KO mice histopathological examination wherein, insulin staining decreases in the mid to late postweaning period. The inability of islets to respond to high glucose is a feature of immature islet cells, specifically β cells. When compared to adult islets, neonatal islets are functionally immature and fail to match the secretory capacity of adult mouse islets in response to high glucose [34]. Functional maturity is coupled to changes in the transcriptional program. With age, mouse islets display increased expression of metabolism genes, encoding for urocortin 3, lactate dehydrogenase, pyruvate carboxylase, malic enzyme, malate dehydrogenase, and others [35]. Given ROS has an establish role in pancreatic development [36], it is plausible that, GSH depletion-induced ROS influences the maturation-state of the islet in *Gclc* KO mice.

Given our focus on newly weaned *Gclc* KO mice, we confirmed that islets had decreased GCLC (1.83% of CTRL) expression and, subsequently reduced GSH biosynthesis (14% of CTRL). We confirmed that, the loss of GCLC conferred oxidative stress by examining the expression levels of established oxidative stress-related genes including: the two major cellular redox systems, GSH (*Gclm, Gsr, Gpx1, and Gpx4*) and thioredoxin (Trx) (*Txn1, Txn2, Txnrd1, and Txnip*) and antioxidant defense (*Nrf2, Hmox1*, *Nqo1*, *Bach1*). Of the two major cellular redox systems, only two genes were significantly changed. *Gclc* expression, as expected, was significantly decreased. Of genes related to antioxidant defense, only *Hmox1* and *Nqo1* were significantly increased in newly weaned *Gclc* KO mouse islets. We did not observe a significant change in *Nrf2* expression. This is unsurprising as, Nrf2 activity is regulated post-transcriptionally [37]. *Hmox1* and *Nqo1* are direct targets of *Nrf2* suggesting *Nrf2* activity. We also observed a significant increase in *Gpx4* expression **(6b)**. The *Gpx4* gene encodes for glutathione peroxidase 4 (Gpx4). Gpx4 is a cytosolic peroxidase which reduces phospholipid hydroperoxides, at the expense of GSH [38]. The GSH-Gpx4 pathway is known to regulate ferroptosis, iron-dependent programmed necrotic cell death caused by lipid peroxidation. At first glance, ferroptosis provides an attractive mechanism for the observed islet dysfunction and eventual, cell death. However, like *Hmox1* and *Nqo1*, *Gpx4* are Nrf2-mediated genes. Nrf2 activity is known to suppress ferroptosis [39]. Additionally, the expression of *Bach1*, a known positive transcriptional regulator of ferroptosis, was unchanged in *Gclc* KO islets when compared to *Gclc* CTRL. This suggests that, the oxidative-stress response in newly weaned islets is mediated by Nrf2 and future experiments should utilize imaging techniques to confirm that, Nrf2 is translocated to the nucleus of islet cells and promoting transcription of the antioxidant response element (ARE) genes.

We observed significantly increased relative islet size in *Gclc* KO mice when compared to *Gclc* CTRL mice at P21. A role of GSH is to prevent the accumulation of ROS and ROS-mediative oxidative stress. Upon GSH depletion ROS increases [40, 41]. A small increase in ROS, particularly H_2_O_2_ can increase proliferation while, a large increase in ROS is pathogenic. β cells included, proliferate in response to modest H_2_O_2_ exposure [42]. Accordingly, we hypothesized that the significant increase in relative islet size in newly weaned *Gclc* KO mice was driven by increased proliferation (i.e., hyperplasia). Analysis of *Mki67* (Ki67) expression, a well-establish marker of proliferation, indicated no increase in proliferation of newly weaned *Gclc* KO mouse islets. Re-examination of histology of the P21 islets revealed hypertrophic islet regions, wherein, portions of the islet were swollen with granular cytoplasm. These finding suggest that the increase in islet size is due to hypertrophy however, nuclear counting experiments should be conducted to confirm.

When characterizing the nature of increased islet size in the newly weaned *Gclc* KO mice, we analyzed the level of expression of cell-cycle related genes, *Cdkn1a* (p21) and *Cdkn2a* (p16). Surprisingly, the expression of *Cdkn1a* (p21) but not, *Cdkn2a* (p16) was significantly higher in the *Gclc* KO mouse islets when compared to *Gclc* CTRL. *Cdkn1a* (p21) and *Cdkn2a* (p16) are genes, encoding for CDKN1A and CDKN2A proteins which, inhibits the activity of cyclin-dependent kinases and arrests cell cycle progression [43]. This observation was unusual given, during the first postnatal month, the islet cells, notably β cells are highly proliferative [44]. Interestingly, the observed cell-cycle inhibition, taken together with, the observed hypertrophic islet regions is suggestion of senescent islet cells. Senescence is a progressive cellular state, that can be driven by DNA damage, oxidative stress, and mitochondrial dysfunction [45, 46]. Early senescence presents with increased p21 expression [47–49]. If the stress continues, the growth arrest is stabilized by the increased expression of p16 and accompanied by distinct changes to cell morphology. This morphology includes, enlarged flattened cells with granularity [50]. Additionally, progressively senescence cells acquire a Senescence-Associated Secretory Phenotype (SASP) and will secrete pro-inflammatory cytokines, like IL-6, TNF-α, thus, engaging the immune system [51]. We sought to better characterize the potential senescent phenotype and, performed gene expression analysis for SASP genes. Interestingly, *Il-6* expression was significantly increased in newly weaned *Gclc* KO mouse islets but, *Tnf-α* was unchanged. This might be explained by histological analysis of the *Gclc* KO mouse islets, which demonstrate that, portions of cells in a single islet might be progressed further along the senescence paradigm. Specifically, the enlarged, granular cells are progressed to late senescence whereas the seemingly “normal” portions of the islet are in early senescence. SASP progressively engages the immune system and immune cells (particularly, macrophages), are recruited to phagocytize senescence cells [52]. The upregulation in *IL-6* promoted us to characterize immune system involvement. During the mid to late weaning period, *Gclc* KO mice displayed increased surveillance by macrophages not, neutrophils, t cells, or b cells **(Fig S3a)**. It should be noted, no single marker of senescent cells exists rather, a collection of markers describes a senescent cell. Additionally, the senescent phenotype can differ by cell type and stage (early to late) of senescence [26, 53]. However, our data suggests that, newly weaned, oxidatively-stressed *Gclc* KO mouse islets may be in a state of early senescence and, the progression of this senescence phenotype might engage the immune system to clear senescent cells and contribute to the severe diabetes phenotype. However, further characterization of the progression of this cell state in our model is warranted.

The present study is, to our knowledge, the first to describe the lack of *Gclc* expression and GSH biosynthesis during pancreatic islet development as a cause for a diabetes phenotype, *in vivo*. These findings enhance our understanding of the natural progression of diabetes and other pancreatic islet pathologies.

## Author Contributions

V.V. and Y.C. conceived of the study. E.A.D., Y.C., S.S., R.C., R.G.K., D.C.T., and V.V., designed the experiments. E.A.D. and Y.W., performed the experiments. E.A.D., Y.C., D.O., B.T., T.F., R.C., R.G.K., C.T.S., D.W.T., D.C.T., and V.V., analyzed data, discussed results, wrote and edited the manuscript.

## Acknowledgements

We would like to acknowledge the Yale Liver Center, for the use of their Olympus BX51, brightfield microscope. The use of Yale Liver Center equipment is supported by award NIH P30 DK034989. We would like to thank the Flavell Laboratory at Yale University for the use of their Keyence, All-in-One Fluorescence Microscope.

---

**Fig. S1a.** *Gclc KO mice display significant weight loss from postnatal day (P) 40 to P42.* Bodyweight of *Gclc* KO mice compared to *Gclc* CTRL mice at postnatal day (P) 42. **a.)** Bodyweight (g). Results are presented as mean ± S.D., n = 3-5 mice/genotype. Statistical analysis was performed using a Two-Way ANOVA with Tukey’s Post Hoc test. Significance was determined at *p-value < 0.05, **p-value < 0.01, ns or no = non-significant.

**Fig. S2a.** *Gclc KO mice display significant hyperglycemia at postnatal day (P) P42.* Fasting blood glucose levels of *Gclc* KO mice compared to *Gclc* CTRL mice at postnatal day (P) 42. **a.)** 6-hr morning fast (∼8:00am – 2:00pm), fasting blood glucose (mg/dL). Results are presented as mean ± S.D., n = 3 mice/genotype. Statistical analysis was performed using an unpaired t-test. Significance was determined at *p-value < 0.05, **p-value < 0.01, ns or no * = non-significant.

**Fig. S3a:** *Gclc KO mice of mid to late weaning age display immune surveillance by islet-specific macrophages.* Representative images of stained pancreata from *Gclc* CTRL and *Gclc* KO mice at postnatal day (P) 35. a.) Immunohistochemical staining for immune cells, macrophage (F4/80), neutrophil (Ly6G), T cell (CD5), and B cell (CD79α).

