## Supplementary Material for "Endocrine pancreas-specific *Gclc* gene deletion causes a severe diabetes phenotype"

**Fig. S1:** *Gclc* KO mice display significant weight loss from postnatal day (P) 40 to P42.

S1a.)

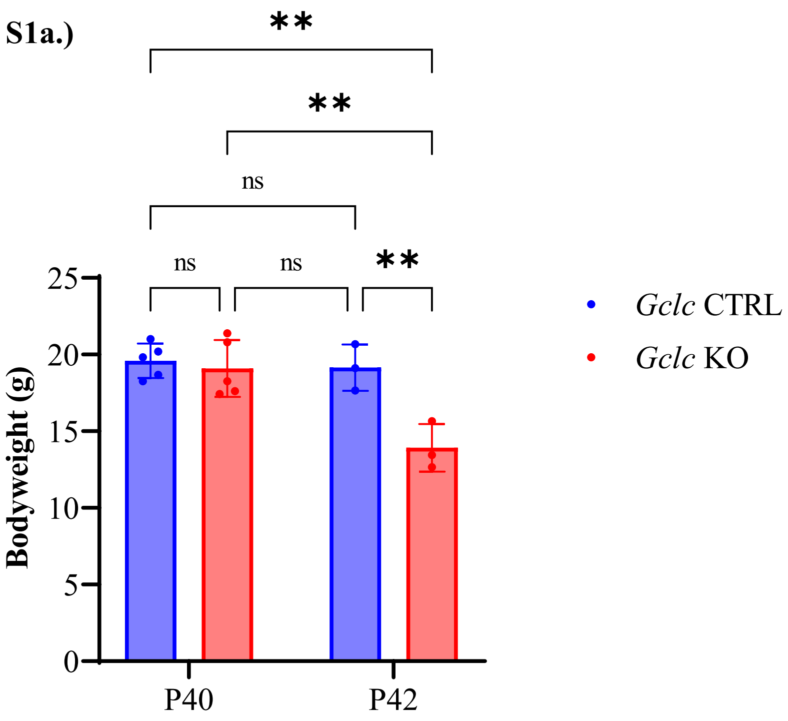


**Fig. S2:** *Gclc* KO mice display significant hyperglycemia at postnatal day (P) P42.

S2a.)

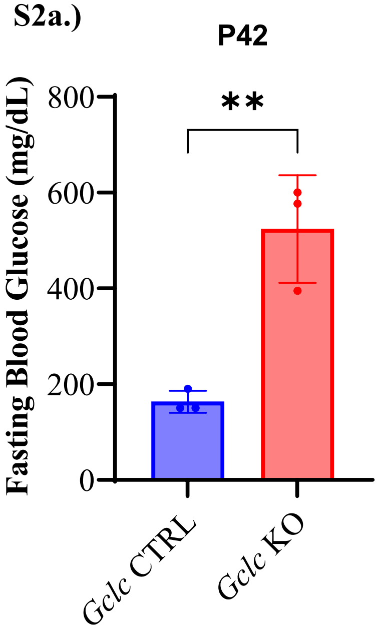

**Fig. S3:** *Gclc KO mice of mid to late weaning age display immune surveillance by islet-specific macrophages.*

**S3a.)**

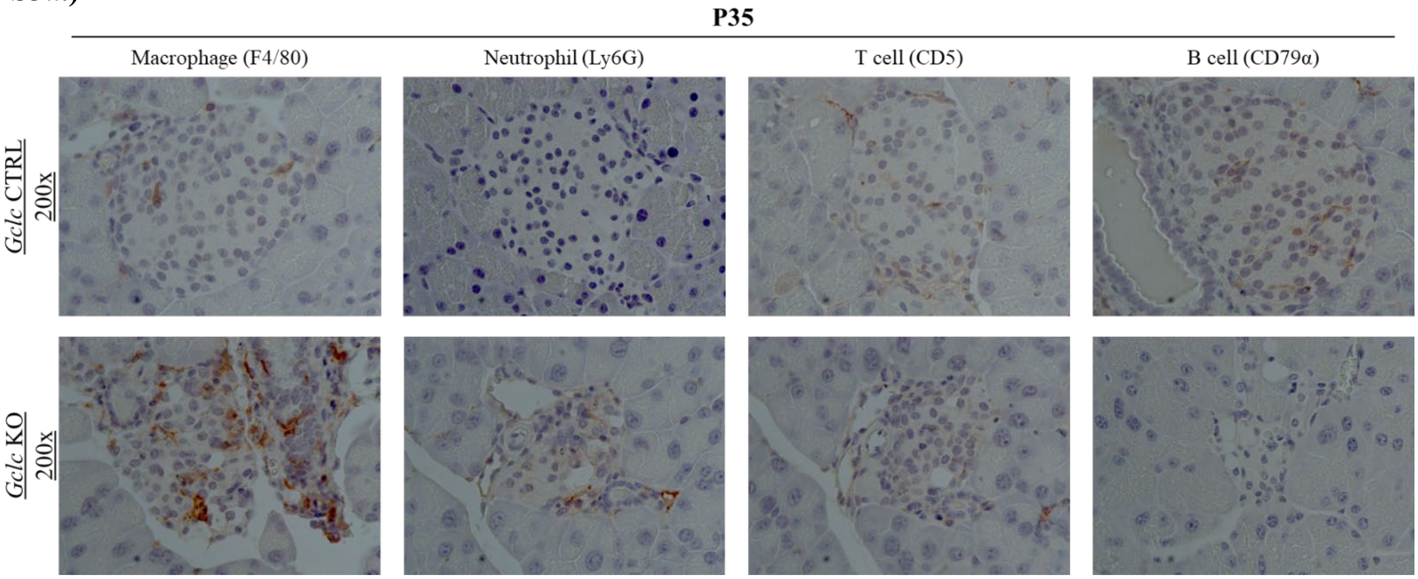

**Supplementary Table S1**. Sequence of primers used for RT-PCR analysis.

| **Gene** | **Forward primer (5'-3')** | **Reverse primer (3'-5')** |
|---|---|---|
| *Bach1* | ATCCATGACATCCGCAGAAG | GAATGTGGTCTCGCTCCTTC |
| *Bach1* | ATCCATGACATCCGCAGAAG | GAATGTGGTCTCGCTCCTTC |
| *Cdkn1a* | TCGCTGTCTTGCACTCTGGTGT | CCAATCTGCGCTTGGAGTGATAG |
| *Cdkn2a* | CTCTGGCTTTCGTGAACATG | TCGAATCTGCACCGTAGTTG |
| *Gcg* | GGATTGCTTATAATGCTGGTGC | TGTGAGTGGCGTTTGTCTTC |
| *Gck* | CTTCACCTTCTCCTTCCCTG | ATCTCAAAGTCCCCTCTCCT |
| *Gclc* | GGCGATGTTCTTGAGACTCTGC | TTCCTTCGATCATGTAACTCCCATA |
| *Gclm* | ACCGGGAACCTGCTCAACT | GCATGGGACATGGTGCATTCC |
| *Glut2* | GTCACTATGCTCTGGTCTCTG | CAAGAGGGCTCCAGTCAATG |
| *Gpx1* | CCACCGTGTATGCCTTCTCC | AGAGAGACGCGACATTCTCAAT |
| *Gpx4* | ATGAAAGTCCAGCCCAAGG | GGTCCTTCTCTATCACCTG |
| *Hmox1* | GCCACCAAGGAGGTACACAT | GCTTGTTGCGCTCTATCTCC |
| *Hnf6* | AGCAAACTCAAGTCGGGTC | GTGTTGCCTCTGTCCTTCC |
| *IL-6* | TGCAAGAGACTTCCATCCAG | TGAAGTCTCCTCTCCGGACT |
| *Ins1* | ATCAGAGACCATCAGCAAGC | GTTTGACAAAAGCCTGGGTG |
| *Ins2* | CGTGGCTTCTTCTACACACCCA | TCCAGTGCCAAGGTCTGAAGGT |
| *MafA* | AGTCGTGCCGCTTCAAG | CGCCAACTTCTCGTATTTCTCC |
| *MafB* | AGAAACATCACCTGGAGAACG | TTCTCGCACTTGACCTTGTAG |
| *Mki67* | GGGTTACTATAGATGAGCCTGTG | CCTGGAAATTCTGTTGGCTTG |
| *Mmp3* | CATAATGATCTCCTTTGCAGTTGG | CATCCTCTGTCCATCGTTCATC |
| *NeuroD* | CTCCAGGGTTATGAGATCGTC | GTCCTGAGAACTGAGACACTC |
| *Ngn3* | CTTTTGAGTCGGGAGAACTAGG | GTATGAGAGTGTGGCTAGGTG |
| *Nkx6.1* | TCTCTGGACAGCAAATCTTCG | CTTGGTCCTGCGGTTCTG |
| *Nq01* | GCGTTCGGTATTACGATCCTC | ACCTCCCATCCTCTCTTCTTC |
| *Nrf2* | CCTCAGCATGATGGACTTGG | CTCATAGTCCTTCTGTCGCTG |
| *Pax6* | TGGTGTCTTTGTCAACGGG | GTTTTCATTGTCCAGCACCTG |
| *Pdx1* | CCCTTTCCCGTGGATGAAATC | GAATTCCTTCTCCAGCTCCAG |
| *Sst* | CCACCGGGAAACAGGAACTG | TTGCTGGGTTCGAGTTGGC |
| *Timp1* | CTCTGGCATCTGGCATCC | TGGTCTCGTTGATTTCTGGG |
| *TnfA* | AAATGGCCTCCCTCTCAT | CCTCCACTTGGTGGTTTG |
| *Txn1* | AATGGTGAAGCTGATCGAGAG | TTGTCACAGAGGGAATGGAAG |
| *Txn2* | AGTTGTCAACAGTGAGACACC | GTGATCGTCAATGTCCACTTTG |
| *Txnip* | GTTGCGTAGACTACTGGGTGAAG | CTCCTTTTTGGCAGACACTGGTG |
| *Txnrd1* | CCAACTTCAAAAGCTGCCAAC | AATCCAAGACCAGCACTTTCT |

**Supplementary Table S2.** Multiple reaction monitoring (MRM) table of transitions for glutathione used for measurements on the Waters Xevo TQ-S-micro triple quadrupole mass spectrometer. Bolded transitions were used for quantification. Non-bolded transitions were used for qualification.

| Metabolite | Precursor ion (m/z) | Product ion (m/z) | Cone voltage (V) | Collision energy (eV) | Dwell time (ms) |
|---|---|---|---|---|---|
| GSH | 307.98 | 179.01 | 40 | 10 | 0.025 |
| **GSH-NEM (Transition 1)** | **433.13** | **303.99** | **42** | **10** | **0.025** |
| GSH-NEM (Transition 2) | 433.13 | 200.93 | 42 | 22 | 0.025 |
